# Sexually dimorphic control of aggression by androgen signaling in a cichlid

**DOI:** 10.1101/2024.04.03.587979

**Authors:** Lillian R. Jackson, Beau Alward

## Abstract

Innate social behaviors like aggression are modulated by sex steroid hormones such as androgens and estrogens. However, we know little about how the same hormone regulates similar behaviors in both sexes. We investigated the role of androgenic signaling in the regulation of aggression in *Astatotilapia burtoni*, a social fish in which males and females perform similar aggressive behaviors. We used ARα knockout (KO) animals for this study, which was recently shown to be required for male-typical aggression and mating. Surprisingly, ARα KO females did not show deficits in aggression. We also determined that females lacking the other AR, ARβ showed normal levels of aggression. Blocking both ARs pharmacologically confirmed that neither AR is necessary for aggression in females. However, ARα KO males showed clear deficits in attacks. Thus, in *A. burtoni* there appears to be a sexual dimorphism in the role of ARα in the control of aggression.

## Introduction

Across many species, social behaviors such as aggression are sexually dimorphic (Nelson and Kriegsfeld, 2017). These sex differences in behavior likely result from sexual selection, in which certain behavioral or morphological features in a competing sex are favored and persist over time (Zahavi and Zahavi, 1999). From a proximate perspective, sex differences in behavior may arise due to the actions of sex steroid hormones such as androgens like testosterone and its androgenic and estrogenic metabolites occurring during early development and/or adulthood (McCarthy, 2016; McCarthy and Arnold, 2011). As described by McCarthy and colleagues (McCarthy et al., 2012) there are multiple types of sex differences including 1) sexual dimorphisms in which a phenotypical endpoint may be substantially more prevalent in one sex and not the other; 2) sex differences in which on average there is a difference in a phenotypical endpoint that spans a continuum between the sexes; and 3) sex convergence where a phenotypical endpoint is similar but the neural mechanisms are different.

Investigations of the causes of the diverse types of sex differences can yield novel insights into the fundamental mechanisms of social behavior more broadly. For example, studying the neural basis of sexually-dimorphic behavior may reveal the level of plasticity within certain brain circuits if a behavior can experimentally be activated in one sex that normally does not perform a behavior, thus providing access to the mechanisms underlying plasticity more broadly (Arnold, 1980; Ball, GF, Balthazart, J, McCarthy, 2014; Madison et al., 2014; McCarthy et al., 2012; McCarthy and Arnold, 2011). Additionally, investigating convergent sex differences in species in which males and females perform the same behavior may provide knowledge of the diverse ways in which distinct upstream molecular and neural mechanisms can converge on identical physical outputs.

Teleost fishes exhibit clear sex differences in reproductive physiology and behavior that are impacted by sex steroid hormones (Godwin, 2010; Tokarz et al., 2015). We recently found in the African cichlid fish *Astatotilapia burtoni*, an organism in which mechanisms of social behavior can be studied in the lab, a dynamic array of sex differences in the performance of aggression. Specifically, during a mirror assay in which *A. burtoni* react to their reflection with aggressive displays, males and females perform both similar and sex-typical behaviors (Jackson et al., 2024). For instance, both males and females perform attacks and lateral displays, but males perform significantly more of the latter. Only males perform rostral displays while only females perform quivers during the mirror assay. Intriguingly, males *do* perform quivers as part of a stereotyped mating ritual with females, indicating females use an identical behavior compared to males but for a different purpose. Finally, male *A. burtoni* with mutant *ar1*, the gene encoding androgen receptor (AR) α, performed significantly fewer lateral displays and quivers during a dyadic social interaction in which they were housed with a stimulus male and three females (Alward et al., 2020).

Given that males and females perform similar and distinct behaviors during an aggression assay, we wondered whether they were governed by the same molecular mechanisms. Specifically, we tested whether female aggression—which includes lateral displays and quivers—is dependent on ARα. As such, we tested wild-type females and females deficient in ARα in a mirror assay. Based on findings in these mutants, we also tested females deficient in ARβ, a second AR present in teleosts, and females treated with an AR pharmacological antagonist. We ultimately discovered that, unlike in male *A. burtoni*, AR signaling is not required for any aggressive behaviors displayed by female *A. burtoni*. Since ARα is not required for quivers in females behaving aggressively towards a mirror, but it is for males performing courtship towards females (Alward et al., 2020), this also implies a convergent sex difference in the control of the same behavior. Our findings have important implications for understanding the hormonal control of social behaviors in teleosts and other species.

## Methods

### Animal Subjects

Adult wild-type *A. burtoni* were bred from a wild-caught stock that originated from Lake Tanganyika, Africa. Adult ARα and ARβ mutant *A. burtoni* were bred from CRISPR/Cas-9 gene-edited lines generated as previously described in Alward et al., 2020. ARα mutations consisted of a frameshift deletion of 50 base pairs (bp) and ARβ mutations possessed a deletion of 5 bp. Wild-type (WT) fish had two functional alleles of the ARs, heterozygous mutants (Het) had one functional allele of either ARα or ARβ, and homozygous mutants (KO) had no functional alleles of either ARα or ARβ. Fish were housed in environmental conditions that mimic their natural equatorial habitat (28 °C; pH 8.0; 12:12 h light/dark cycle with full spectrum illumination; constant aeration). Aquaria contained gravel-covered bottoms with terra cotta pots cut in half to serve as shelters. Fish were fed cichlid flakes daily. All experimental procedures were approved by the University of Houston Institutional Animal Care and Use Committee (Protocol #202000001).

### Mirror assay for assessing aggression in AR mutant A. burtoni

As in recent work, female aggression was assessed using a mirror assay. ARα (WT n=11, Het n=13, KO n=7) and ARβ (WT n=7, Het n=11, KO n=9) mutant females were housed in a pre-assay tank for approximately two weeks prior to being assayed in a mirror assay as described in Jackson et al., 2024. The pre-assay tank consisted of a 16-gallon plexiglass tank with a clear perforated divider that separated the focal females from a wild-type stimulus male on separate halves. Subjects were housed in groups of 3 females at a time. Female genotypes were unknown while housed, but ARα and ARβ females were separately housed in pre-assay tanks. After two weeks in the pre-assay tank, each subject’s behaviors were recorded in a mirror assay. The mirror assay, described in Jackson et al., 2024, consisted of a white tub (Sterilite; 400 mm x 318 mm x 12 mm) filled 6 cm deep with UV-sterilized aquaria water with a mirror replacing one side of the assay tank. An opaque cover was first placed on the mirror for a 15-minute habituation period to allow subjects to acclimate to the assay tank. The opaque cover was then removed, and 30 minutes of behavior was recorded. Immediately after the recording stopped, a fin clip sample was collected from each subject to determine their respective AR genotype (See “Genotyping”).

A series of mirror assays using the same approach as above was also performed in ARα mutant males (WT n=8, KO n=10) to confirm that ARα KO males possess similar deficits in behaviors in a mirror assay as they did in recently published work that used a different behavioral assay (Alward et al., 2020). Immediately after the conclusion of the assay recording, a fin clip was collected from each subject for subsequent genotyping (See “Genotyping”).

### Genotyping

Immediately following the behavioral assay, each subject was fin-clipped for genotyping. Using scissors cleaned with ethanol, a 1-to 2-mm section of the caudal fin was excised and placed into an individual PCR tube. To extract DNA, 180 μL of NaOH (50 mM) was added to the DNA sample. The sample was incubated at 94°C for 15 minutes. After this incubation, 20 μL of Tris HCl (pH = 8) was added to the sample and then vortexed and spun down using a minicentrifuge for ∼ 5 seconds. Samples were placed at -20°C for at least 15 minutes before PCR amplification of mutated regions of ARα or ARβ.

### CA injections and behavior recording

Gravid wild-type *A. burtoni* females were removed from a community tank and injected intraperitoneally with either a sesame oil vehicle or cyproterone acetate (CA, concentration = 0.83 mg/gbw). CA injected (n = 9) and sesame oil vehicle (SV) (n = 10) injected females were each placed into a 20.8 L glass tank containing gravel and constant aeration and were observed for 30 minutes following the injection to monitor recovery. Two stimulus females were added to each tank with an injected female after the injection recovery. 2 days after the injection, focal injected females were assayed in a mirror assay as described above.

### Behavioral analysis

Behavior was recorded using a digital video camera and was quantified using the freely available BORIS (Behavioral Observation Research Interactive Software). Multiple types of behavior were quantified: behaviors considered not to be aggressive or “neutral” (tap, graze); subordinate behaviors (flee, retreat); aggressive behaviors (attack, lateral display, quiver, rostral display). Additionally, a female reproductive behavior (egg release) was included after two ARα females released and picked up their eggs during the mirror assay trials (one WT and one Het female). Descriptions of these behaviors are included in Table 1.

**Table 1.** Ethogram used to score behaviors in BORIS.

| **Behavior** | **Description** | **Category** |
| --- | --- | --- |
| Tap | Fish makes gentle mouth contact with the mirror | Neutral |
| Graze | Fish swims alongside the mirror for at least 3 seconds | Neutral |
| Flee | Fish makes an abrupt turn and swim away from the mirror | Subordinate |
| Retreat | Fish rapidly retreats from the mirror | Subordinate |
| Attack | Fish rapidly swims towards the mirror, making mouth contact, and vigorously wiggling of the body | Aggressive |
| Lateral Displays | Fish presents the side of the body towards the mirror, conforming the body at times in a slightly convex shape while lateral to the mirror with erect fins | Aggressive |
| Lateral Display Type 1 | More vigorous lateral display | Aggressive |
| Lateral Display Type 2 | Less vigorous lateral display | Aggressive |
| Rostral Display | Fish flares the opercula towards the mirror without making direct contact with the mirror | Aggressive |
| Quiver | Fish makes a rapid vibration of the body towards the mirror while conforming in a convex shape | Aggressive |
| S-curve Display | Fish makes a curved body conformation facing the mirror at a distance |  |
| Egg Release | Female releases eggs and picks them up to hold in her mouth | Reproductive |

Tracking and activity data was collected on pre-mirror and mirror assay videos using Noldus EthoVisionXT software. Center-point detection was used to track the subjects throughout the assay tank. Distance moved (cm) was calculated as the accumulated distance the subject moved throughout the assay duration. Velocity (cm/s) was calculated as distance over time. Activity was measured as the average % of pixels changed over time. We defined a zone close to the mirror to measure the proximity of subjects near the mirror, extending 2.4 cm out from the mirror. Zone duration (s) was calculated as the cumulative duration of time spent in the zone. Zone frequency was calculated as the number of times the center-point of the subject entered the zone. Zone latency was defined as the lapse of time until the zone is first entered. Total and mean distances to the zone (cm) were calculated as the cumulative and mean distances from the zone respectively.

### Morphological analysis

Subjects were assessed for standard length (SL), body mass (BM), gonad mass, and gonadosomatic index [GSI = (gonad mass / body mass) * 100].

### Statistical Analysis

All statistical tests were performed using Graphpad Prism version 10. One-way ANOVAs were used to compare aggressive behaviors between the genotypes for ARα and ARβ mutants. If assumptions were not met, the data were log transformed followed by a one-way ANOVA. If the log transformed data did not correct the data to meet the assumptions, a non-parametric Kruskal-Wallis test was performed. Since none of the main effects for the one-way ANOVAs were significant, no multiple comparisons were performed. To compare aggressive behaviors between WT and KO ARα males, we performed unpaired t-tests. When normality and equality of variance assumptions were not met, a log transformation of the data was performed before conducting an unpaired t-test and if this transformation did not correct the data to meet assumptions, we performed a Mann-Whitney test. Unpaired t-tests were also used to analyze the effects of treatment in the CA experiment.

## Results

### Neither ARα or ARβ are required for aggressive behaviors during a mirror assay in female A. burtoni

To determine if female aggression was AR dependent, we assayed ARα and ARβ mutant females in a mirror assay and tested whether AR manipulation affected aggressive behaviors in females like it does in males (Figure 3). We found that ARα wild-type (WT), heterozygous mutants (Het), and homozygous null mutants (KO) did not differ in the number of attacks, lateral displays, and quivers performed in a mirror assay (Figure 1a; attacks, Kruskal Wallis, *p* = 0.8925, H = 0.2274; Table 2; lateral displays, quivers, One-way ANOVAs of log transformation). We also found that ARβ WT, Het, and KO mutants did not differ in the number of attacks, lateral displays, and quivers performed in a mirror assay (Table 2; attacks, lateral displays, quivers, One-way ANOVAs of log transformation).

**Table 2.** Effect of genotype on aggressive behaviors in ARα and ARβ mutant females. Degrees of freedom are written below each F statistic.

| | $AR\alpha$ | | $AR\beta$ | |
| --- | --- | --- | --- | --- |
|  | Test Statistic | P | Test Statistic | P |
| Attacks | H = 0.2274 | 0.8925 | F = 0.5298<br>2, 24 | 0.5954 |
| Lateral Displays | F = 0.08659<br>2, 20 | 0.9174 | F = 0.01824<br>2, 16 | 0.9819 |
| Quivers | F = 0.2534<br>2, 19 | 0.7787 | F = 0.7742<br>2, 10 | 0.4868 |

**Figure 1.**
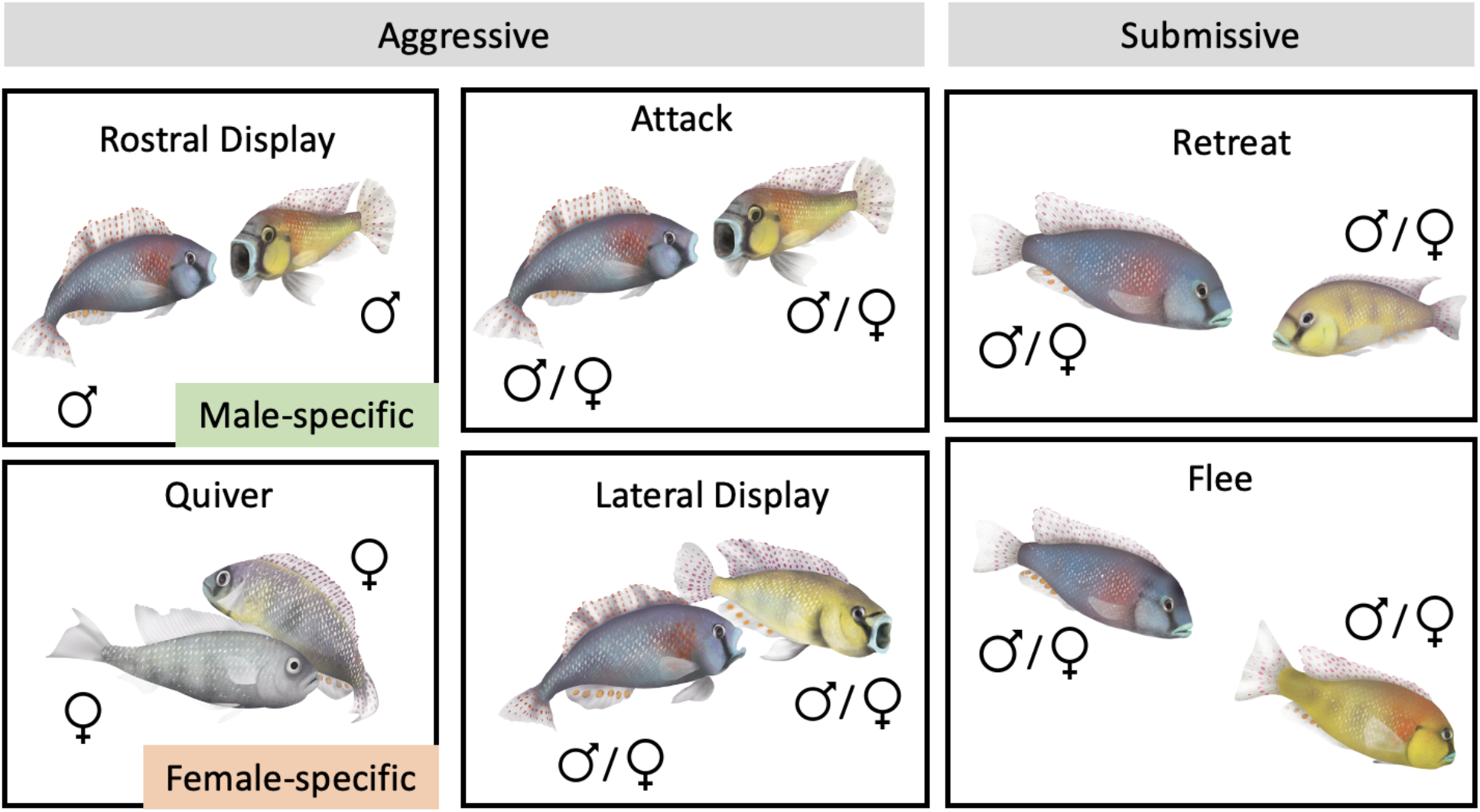
Typical behaviors characterized in a mirror assay in *A. burtoni*. Rostral displays are a male-specific aggressive behavior, and quivers are a female-specific aggressive behavior. Other behaviors are performed by both males and females.

**Figure 2.**
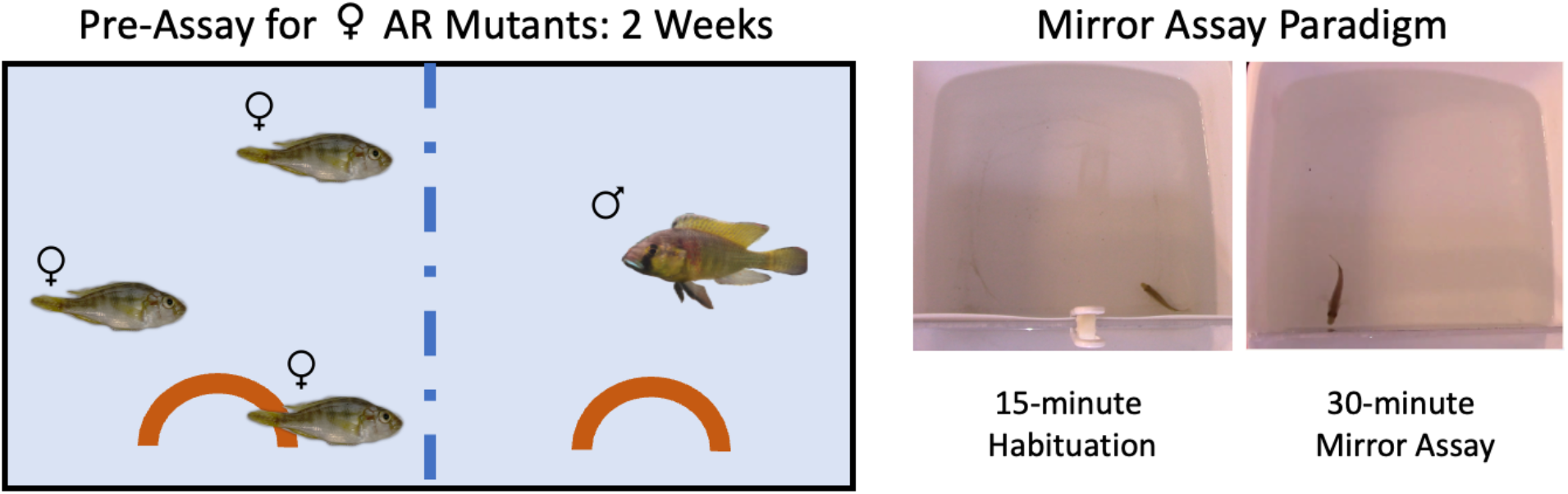
Pre-assay schematic for ARα and ARβ mutant females. Mutant females were housed in a divided tank with a stimulus WT male for 2 weeks and then assayed in a mirror assay.

**Figure 3.**
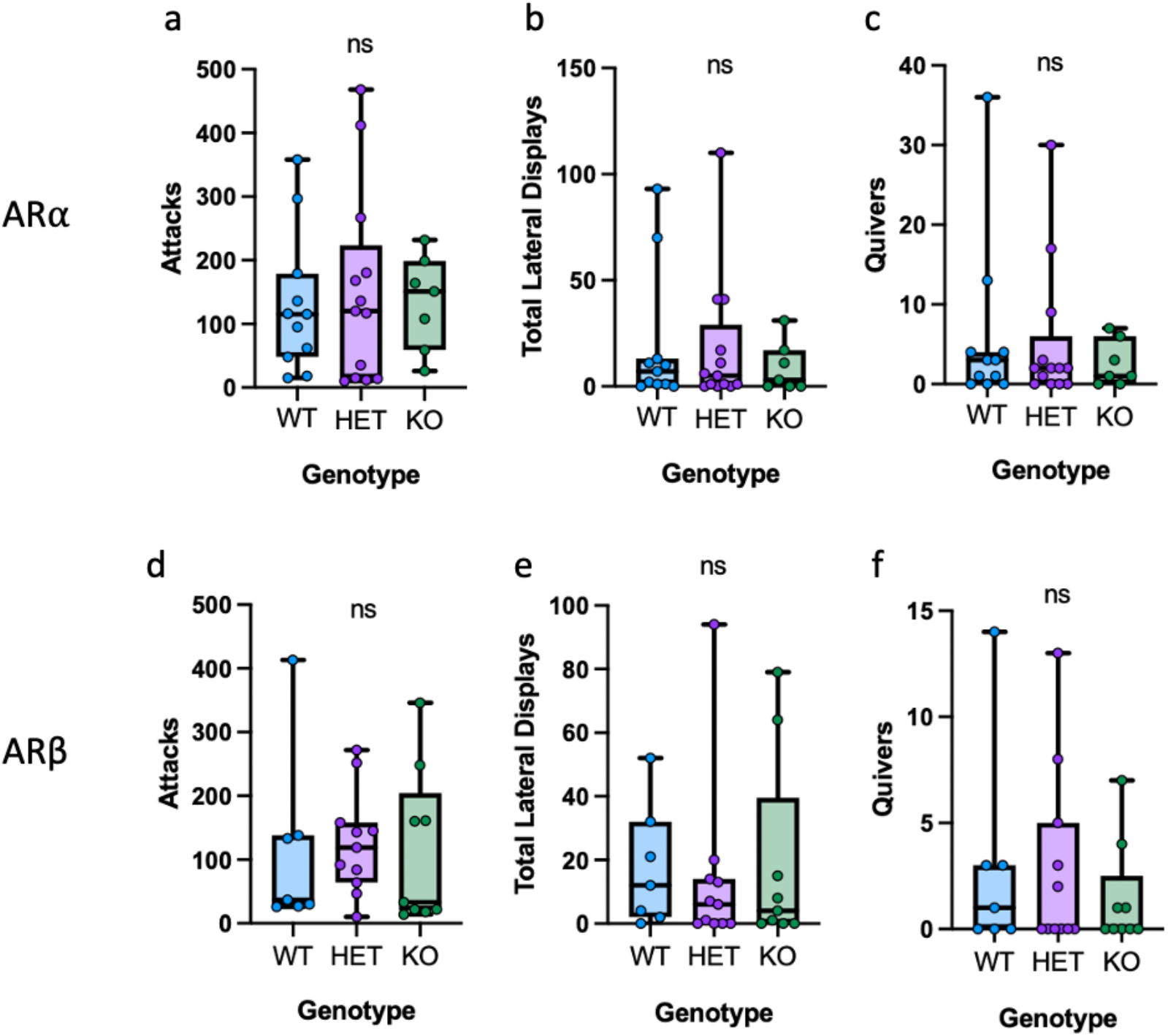
Aggressive behaviors in *A. burtoni* females are not modulated by androgen receptors (ARs). (a)(b)(c) There was no effect of ARα genotype on attacks, lateral displays, and quivers in females. (d)(e)(f) There was no effect of ARβ genotype on attacks, lateral displays, and quivers in females. WT, wild-type; HET, heterozygous mutant; KO, homozygous mutants. ns, not statistically significant.

### ARα is required for aggressive behavior during a mirror assay in males

Since ARα is required for male dominance behaviors (Alward et al., 2020) in which KO males show deficits in quivers and lateral displays, we wanted to confirm that ARα KO males possess deficits in male-typical aggressive behaviors in a mirror assay, ensuring that the results we found in female AR mutants are not due to specifically to the nature of the assay itself. We assayed ARα WT and KO males in a mirror assay and found that ARα KO males show deficits in attacks (Figure 4a; Mann-Whitney, *p* = 0.0452, U = 17.5). WT and KO males did not differ in lateral displays (Figure 4b; Mann-Whitney, *p =* 0.2164, U = 31.5). WT males on average performed more rostral displays but this did not each statistical significance (Figure 4c; Mann-Whitney, *p* = 0.0686, U = 22.5). Only 1/10 ARα KO males performed a rostral display compared to 4/8 WT males. A χ^2^ test comparing these frequencies, however, was not statistically significant (*p=* 0.0597). Males do not perform quivers, a male-typical courtship behavior, during aggressive interactions so it is impossible to assay the necessity of ARα on quivers in males during a mirror assay.

**Figure 4.**
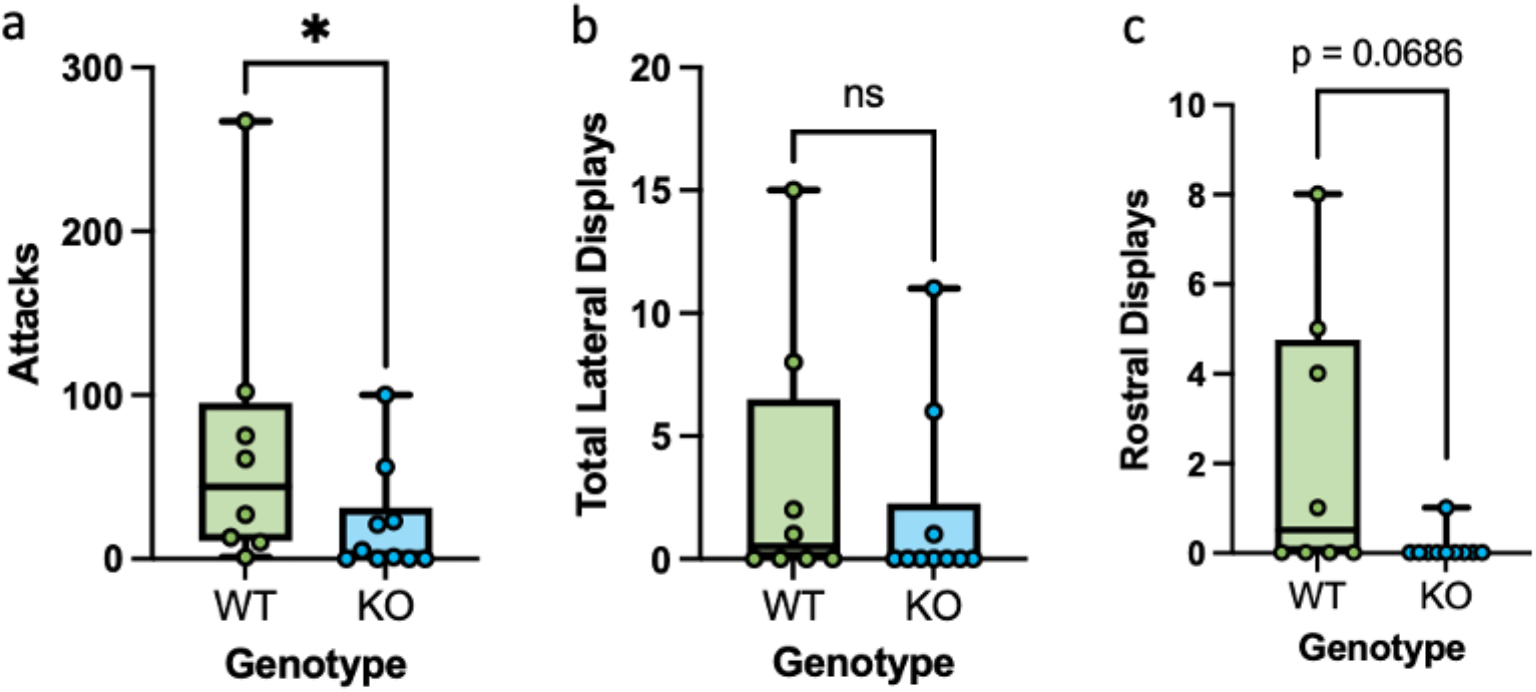
Aggressive behaviors in *A. burtoni* males are modulated by androgen receptor α. (a) WT males performed more attacks than KO males. (b) WT and KO males did not differ in lateral displays. (c) WT males performed more rostral displays on average than KO males, although this was not statistically significant.

### Pharmacological antagonism of ARs does not alter female aggressive behaviors

One interpretation of null findings in ARα and ARβ mutants is that both receptors function redundantly in the control of female aggression during a mirror assay. To test this hypothesis, we injected females before the mirror assay with the steroidal AR antagonist CA. CA has been used in multiple studies aimed at inhibiting both ARs present in teleost fishes, including in *A. burtoni* and other cichlid fish (Alward et al., 2019; O’Connell and Hofmann, 2012; van Breukelen, 2013). We found that females treated with CA do not differ from those injected with vehicle in the number of attacks, lateral displays, and quivers performed in a mirror assay (Figure 5; Unpaired t-tests of log transformation, attacks: *p* = 0.5550, df =16; lateral displays: *p* = 0.4185, df = 11; quivers: *p* = 0.3727, df = 13).

**Figure 5.**
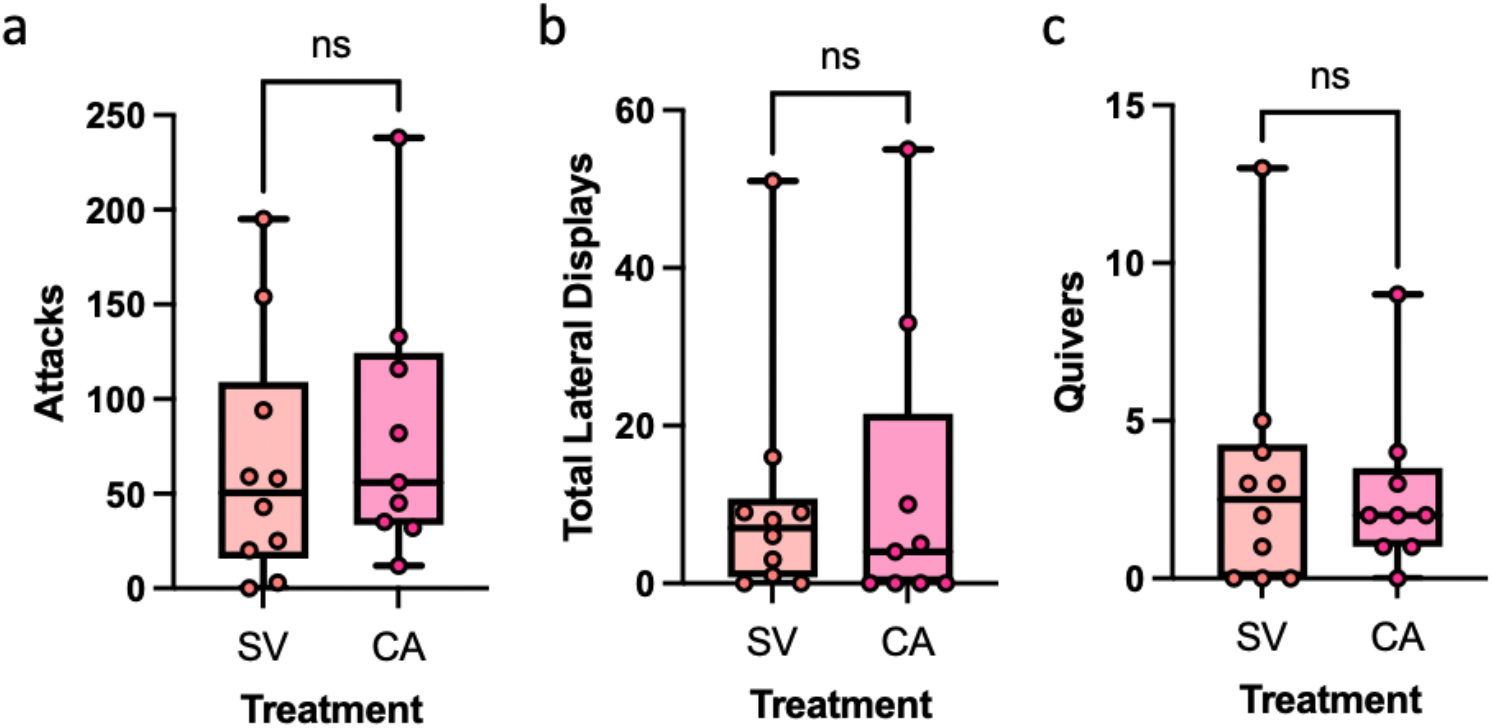
Pharmacological antagonism of AR does not decrease aggressive behaviors in *A. burtoni* females. (a)(b)(c) There was no effect of AR antagonism on attacks, lateral displays, and quivers in females. SV, sesame vehicle; CA, cyproterone acetate. ns, not statistically significant.

## Discussion

In male *A. burtoni*, aggressive and reproductive behaviors are dependent on androgen signaling, specifically through androgen action at ARα (Alward et al., 2020). Previously, we found that females perform similar aggressive behaviors to males in the same behavioral assay and we discovered that females perform a stereotypically identical male-typical reproductive behavior, quivers, in this aggressive context (Jackson et al., 2024). We investigated whether androgen signaling regulates aggressive behaviors in *A. burtoni* females as it does in males. We found that genetic and pharmacological manipulation of androgen receptors in females had no impact on aggressive behaviors, including quivers. ARα mutant males in the mirror assay, however, showed deficits in aggressive behavior, like in recently published work (Alward et al., 2020). These results suggest a sexually dimorphic role of ARα in the regulation of aggression in *A. burtoni*.

The dependence of aggressive behavior on androgen signaling is not unique to *A. burtoni* males. AR signaling also regulates male-typical behavioral displays and aggression in male mice, rats, birds, lizards, and fish. *AR* gene inactivation in male mice causes a severe reduction in aggressive behaviors (Sato et al., 2004), and male mice lacking AR in the nervous system were less aggressive than controls, spending less time fighting and had a longer interval between successive attacks (Juntti et al., 2010). In male song sparrows, administration of the antiandrogen flutamide decreased some aggressive behaviors during the pre-breeding season in freely living birds as well as laboratory-held birds under long-day (LD) conditions (Sperry et al., 2010). The AR antagonist flutamide also reduced aggressive behavior in male red-winged blackbirds (*Agelaius phoeniceus)* and male European robins (*Erithacus rubecula*) (Schwabl and Kriner, 1991; Searcy and Wingfield, 1980). In *Anolis sagrei*, CA decreased male territorial aggression (Tokarz, 1987). CA also decreased the number of aggressive displays in convict cichlid (*Amatitlania nigrofasciata*) males in a social challenge context (Sessa et al., 2013). However, despite the evidence that male aggression is influenced by androgen signaling, it is important to note that these studies did not include females in their analysis of the role of AR in aggression. In many species, females are much less aggressive than males, making it difficult to assess their aggressivity. In resident-intruder assays in mice, male but not female residents attacks intruders in their home cage (Miczek et al., 2001). Our study is unique in that both males and females perform aggressive behaviors, which allowed us to identify the molecular mechanisms that contribute to the same behaviors in each sex.

Aggressive behaviors are first initiated by the assessment of an aggression-provoking cue through the sensory modalities (Lischinsky and Lin, 2020). For some animals, the stimuli that provoke aggression are conveyed through one sensory modality, while other animals integrate signals received through several sensory modalities. For example, rodents rely mainly on olfactory cues to detect aggression-provoking stimuli (Dulac and Torello, 2003) while songbirds rely on auditory input (Searcy et al., 2006). For animals whose aggression is triggered by specific sensory signals, these aggression-provoking stimuli are likely transmitted through a species-specific cue detection circuit, directing information to the core aggression circuit as proposed in (Lischinsky and Lin, 2020). This cue detection circuit takes sensory cues, for example like olfactory signals in rodents, directs them to neural circuitry that determines whether a behavioral output is triggered or not. The core aggression circuit (CAC) is comprised of the medial amygdala, bed nucleus of the stria terminalis, ventrolateral part of the ventromedial hypothalamus, and the ventral part of the premammilary nucleus (Lischinsky and Lin, 2020). The CAC triggers motor output in response to the initial aggression-provoking stimuli, resulting in the production of innate and stereotypical behaviors. Here is where we identify sexually dimorphic aggressive behaviors, caused by sex differences in the aggression circuits. In rodents, the aggression circuitry is shared between the sexes and is qualitatively similar in males and females (Hashikawa et al., 2018). Despite these similarities though, sexually dimorphic aggressive behaviors have been characterized in rodents. Therefore, these quantitative differences in aggressive behaviors may be due to the sex difference in the number of the aggression-related cells within each aggression relay or the sex difference in response magnitude of these cells (Hashikawa et al., 2018). Finally, the molecular mechanisms of the same behaviors may be sexually dimorphic – as we are proposing here with role of androgen signaling in mediating aggression in male, but not female *A. burtoni*. In our previous research, we found that the anterior tuberal nucleus (ATn, the putative VMH homolog in fish) is functionally connected to the preoptic area (POA) in males, but not females after exposure to an aggressive context (Jackson et al 2024). Importantly, both the ATN and POA express ARα in *A. burtoni* (Harbott et al., 2007; Lopez and Alward, 2023). This steroid-sensitive, sexually dimorphic connection could be a potential upstream circuit linking the CAC to behavioral output, explaining the sex differences in aggressive behaviors triggered by an identical context.

We found deficits in aggressive behaviors in male ARα mutants in a mirror assay, in which KO males perform fewer attacks than WT males. Other behavioral deficits were shown in Alward et al., 2020 in which ARα KO males do not perform lateral displays. Although we did not find the same effect on lateral displays in our assay as was found in previous work (Alward et al., 2020), there are important differences between studies to consider. For example, the social context here (a mirror assay) is different compared to the previous work (a dyadic social interaction, where a focal male was housed with a smaller stimulus male and three females). Therefore, the different behavioral deficits may have to do with distinct behavioral strategies that are employed in different social contexts. Indeed, in the mirror assay, males are size matched with their reflection and may adjust their behavior accordingly towards their “opponents”. Additionally, the “resident effect” from a resident-intruder interaction, as described in Alward et al., 2020 and by many others (Nijman and Heuts, 2011; Owen and Gordon, 2005), is removed in a mirror assay with a limited territory size. In the study finding the deficits in lateral displays, the focal males had a significant size advantage over their opponents (Alward et al., 2020). This may also indicate the intriguing possibility that the role of androgen signaling in modulating aggression in males differs based on social context. It is not unprecedented for a steroid-dependent behavior to be impacted differently based on experience/social context. In solitary male mice, the activation of PR+ VMHvl neurons are necessary and sufficient for aggression (Yang et al., 2017). However, social context can alter the initiation of aggression with stimulation of PR+ VMHvl neurons in which the initiation of aggression in socially housed males is suppressed (Yang et al., 2017). In our study, perhaps the resident male’s perception of their opponent’s size or their territory alters which aggressive behaviors dependent on androgens are to be performed. These questions can be explored in the future.

If androgen signaling does not mediate female aggression, then what does? we speculate that estrogenic signaling may be involved in regulating aggressive behaviors in female *A. burtoni*. For example, when dominant male *A. burtoni* are treated with estradiol, there is an increase in aggression, namely chases, performed (O’Connell and Hofmann, 2012). Additionally, treatment with an estrogen antagonist, ICI182780, decreased chases performed by dominant males towards other males (O’Connell and Hofmann, 2012). Furthermore, in many species, including mice, rats, and birds there is support for the involvement of estrogen signaling in female aggression. In studies with female mice against female opponents, estrogen receptor α (ERα) KO mice exhibited higher levels of aggression than their WT littermates (Ogawa et al., 1998, 1996). Additionally, ERβ KO females exhibited an increase in postpartum maternal aggression and testosterone-induced aggression towards an olfactory bulbectomized male intruder (Ogawa, 2004). Single acute activation of ERα increased dominance-related agonistic behaviors in ovariectomized (OVX) female mice. Furthermore, acute activation of ERβ increased agonistic behavior in gonadally intact female mice (Clipperton Allen et al., 2010; Clipperton-Allen et al., 2011). In female rats, estradiol benzoate (EB) treatment is ineffective in activating aggression alone, but when combined with dihydrotestosterone propionate (DHTP) is highly effective in the activation of aggression, implicating both estrogenic and androgenic metabolites of testosterone in promoting female aggressive behavior (Van De Poll et al., 1986). Interestingly, even a single injection of estrogens on the day of birth increased agonistic dominance behaviors in adult female rats (Berretti et al., 2014). Finally, physical aggression is shown to be increased by acute estradiol (E2) administration in song sparrows and white-throated sparrows (Heimovics et al., 2015; Merritt et al., 2018), while vocal aggression can be induced by chronic administration of E2 in female white-throated sparrows (Maney et al., 2009). Across many species, there is compelling evidence that estrogen signaling is involved in the production of aggressive behaviors. Therefore, estrogen signaling may modulate aggression in female *A. burtoni* instead of, or in combination with, androgen signaling. Clearly, based on numerous previous studies and the results we have shown, the control of aggressive behaviors by sex steroid hormones across species is dynamic and complex based on multiple factors such as social context and sex. Teleosts like *A. burtoni* that show diverse social behavior differences and similarities as a function of sex will continue to be especially useful in disentangling the molecular and neural mechanisms that underlie innate social behaviors such as aggression.

## Acknowledgements

The authors thank Mariam Dumitrascu and Mariana Lopez for procedural assistance.

## Funding

This research was supported by a Beckman Young Investigator Award from the Arnold and Mabel Beckman Foundation, an NIH grant R35GM142799, and a University of Houston-National Research University Fund startup R0503962 to B.A.A.

## Notes

### Competing Interest Statement

The authors have declared no competing interest.

